# Comparative analysis of perennial and annual *Phaseolus* seed nutrient concentrations

**DOI:** 10.1101/612010

**Authors:** Heather E. Schier, Kathrin A. Eliot, Sterling A. Herron, Lauren K. Landfried, Zoë Migicovsky, Matthew J. Rubin, Allison J. Miller

## Abstract

Malnutrition is a global public health concern and identifying mechanisms to elevate the nutrient output of crops may minimize nutrient deficiencies. Perennial grains within an agroforestry context offers one solution. The development and integration of perennial crops for food has critically influenced dialogue on the ecological intensification of agriculture and agroforestry. However, the nutrient compositions of wild, perennial, herbaceous species, such as those related to the common bean (*Phaseolus vulgaris*) are not well known. In this study, seed amino acid and ion concentrations of perennial and annual *Phaseolus* species were quantified using ionomics and mass spectrometry. No statistical difference was observed for Zn, toxic ions (e.g. As) or essential amino acid concentrations (except threonine) between perennial and annual *Phaseolus* species. However, differences were observed for some nutritionally important ions among and within lifespan groups. Ca, Cu, Fe, Mg, Mn, and P concentrations were higher in annual species. Intraspecific variability in ion concentrations and amino acids was observed within species; further, ion concentrations and amino acids differ among annual species and among perennial species. Ion and amino acid concentration appear to be largely independent of each other. These results suggest variability in ion and amino acid concentrations exist in nature. As new crop candidates are considered for ecological services, nutritional quality should be optimized to maximize nutrient output of sustainable food crops.

## INTRODUCTION

Balancing the nutrition needs of a growing human population with agricultural sustainability is a fundamental challenge for the 21st century. Currently, global nutrition targets set by the World Health Organization are not being met [1] with mineral nutrient and protein energy deficiencies disproportionately impacting lower income countries [2]–[4]. At the same time, demands on agricultural resources (e.g., land, water, soil health) are rising, and solutions to produce nutritious food equitably and sustainably are needed. Is it possible to improve crop nutrition while concomitantly working towards more sustainable agricultural systems? Ongoing efforts explore a variety of ways in which our food system can provide for global nutrition in an ecologically sustainable manner.

One proposed mechanism for improving agriculture sustainability is agroforestry, an agricultural system that incorporates trees, and includes forest farming, alley cropping, silvopasture, riparian buffers, and windbreaks [5]. Agroforestry integrates agriculture and forestry with the goal of enhancing food production while simultaneously improving soil, water, and habitat for biodiversity [6], [7]. It is a form of ecological intensification, a process that harnesses ecological services to sustain production, support ecosystems, and minimize anthropogenic effects of agriculture [8]. As agroforestry research expands [7], [9], attention is focusing in part on the role of herbaceous perennial crops to improve agriculture sustainability [10], [11]. Perennial plants with deep, robust root systems prevent erosion, sequester carbon, and absorb and retain more water [12], [13]; they access and absorb minerals housed deep in the soil, incorporating them into the plant body [14]. Moreover, growing diverse species with variable root architectures (shallow to deep-rooted species) simultaneously may improve the efficiency of nutrient uptake by collectively accessing multiple pools of resources at different soil depths. Agriculturally and ecologically important plant families such as the legume family (Fabaceae) contain thousands of perennial, herbaceous species that might be strong candidates for use in agroforestry systems [15]. However, natural variation in nutritional qualities of many perennial species, and how nutrition in perennial plants compares to annual relatives, is not well-understood.

Developing nutritious, high-yielding crops is an important consideration for alternative agriculture strategies, including agroforestry. Many nutrients required for core biological processes, including minerals and amino acids, must be acquired through dietary consumption [16]. There are 16 essential mineral nutrients, some of which are required in larger amounts (termed ‘macro’ minerals such as calcium, potassium, magnesium), and others that are required in smaller quantities (trace elements including iron, selenium, zinc). Of the 20 amino acids, nine must be acquired through dietary intake (essential amino acids) and the remaining 11 can be synthesized by the human body (non-essential amino acids). Mineral nutrients and essential amino acids can be acquired through plant-based products, or through a balanced diet that includes animal and plant sources. Quantifying concentrations of essential nutrients and understanding their patterns of covariation in plant species used for food is critical for developing food systems that minimize nutrient deficiencies.

In considering the development of crops for agricultural systems such as agroforestry, attention is focusing on species that provide high nutritional content [7], [9] as well as ecological functionality. Mineral and amino acid concentrations of plant species are influenced by both genetic and environmental factors [17]–[28]. In the common bean, *Phaseolus vulgaris*, uptake of iron, zinc, and amino acids varied across genotypes [18], [20]. Environmental conditions (e.g., production site, soil conditions, temperature patterns) and agricultural practices (irrigation, fertilizer impacts) can also influence nutritive properties of beans [19], [21]–[24], [29]. Many other factors may also influence nutrient quality including position of the seed pod along the plant stem, developmental stage, post-harvest processing, and quantity of anti-nutrients (compounds that interfere with nutrient absorption), such as trypsin inhibitors, hemagglutinin, tannin and phytic acid [18], [19], [29]–[31].

As researchers investigate plant species for crop development and potential incorporation into agroforestry systems, an emerging question is the extent to which mineral and amino acid concentrations vary based on how long plants live. There is some indication that the nutrient composition of seeds may vary depending on lifespan (annual or perennial), and that deeper rooted, long-lived perennials are able to uptake higher amounts of ions from the increased surface area and depths of soil [12], [32], [33]. For example, seeds of perennial wheat varieties were 24-40% higher in copper, iron, manganese and zinc, and 3.5-4.5% higher in protein content compared to annual cultivars [34]. As scientists consider different species for agroforestry and other agricultural systems, an important first step is assessing mineral and amino acid concentrations in candidate plant species, and understanding factors underlying nutrient compositions of these species.

In this study we investigate nutritional qualities of some members of the legume family. Legume species are important components of nearly all agricultural systems, and at least five species of the common bean genus *Phaseolus* have been domesticated in different agricultural centers [35]. The common bean (*P. vulgaris*) was domesticated twice in the Americas [36] and is a major food staple among communities in eastern Africa, Latin America, and Asia, where it provides an excellent source of iron, potassium, selenium and zinc as well as protein [37], [38]. Legumes, including *P. vulgaris,* are of particular interest for agriculture in lower income countries where the magnitude and frequency of deficiencies in mineral nutrients and protein is a public health concern. However, legume production has not kept pace with population growth or production of other commodities, such as cereal grains [39], [40]. Further, *P. vulgaris* is an annual crop sensitive to abiotic stress, including drought and high temperatures, and may be vulnerable to predicted future environmental conditions [41]–[43]. For example, the global economic yield of *P. vulgaris* may be reduced by as much as 58-87% in response to drought [44]. Many wild *Phaseolus* species are underexplored and could serve as a partial replacement or complement to existing *P. vulgaris* agriculture [45].

The incorporation of legumes into agroforestry systems is a primary goal because legumes enhance nutrition and simultaneously provide ecosystem benefits through nitrogen fixation [39], [46], [47]. Broadly, protein content of legumes can range from 12 to 55%, and *Phaseolus* species tend to contain between 20 and 35% protein [47]–[49]. An intraspecific study of *P. vulgaris* found protein content varied from 22 to 39% across 87 accessions [50]. Studies have quantified nutrient composition in some perennial bean species [51], [52], however to our knowledge, differences in mineral nutrient and amino acid concentrations of the seeds of perennial and annual species have not been explored in the legume family.

Here, we compared ion and amino acid concentrations in *Phaseolus* seeds of four perennial species and two annual species in order to 1) assess differences in ion and amino acid concentrations across annual and perennial plant species; 2) quantify natural variation in ion and amino acid concentrations within six wild *Phaseolus* species; and 3) identify correlations between mineral and amino acid concentrations across *Phaseolus* species. Our results expand current understanding of variation and patterns of nutritional qualities of different *Phaseolus* species.

## METHODS

### Study system

*Phaseolus* has been cultivated since ~2500 BP and includes approximately 70 herbaceous species of which at least 18 are perennial [15], [53]–[58]. For this study, we selected four perennial *Phaseolus* species (*P. angustissimus, P. filiformis, P. maculatus, P. polyanthus)* and two annual species (*P. acutifolius, P. vulgaris)* (Table 1). To examine within-species variation, two varieties of *P. acutifolius* were included as a case study. A second case study assessed variation between cultivated and wild accessions of *P. polyanthus*.

**Table 1.**
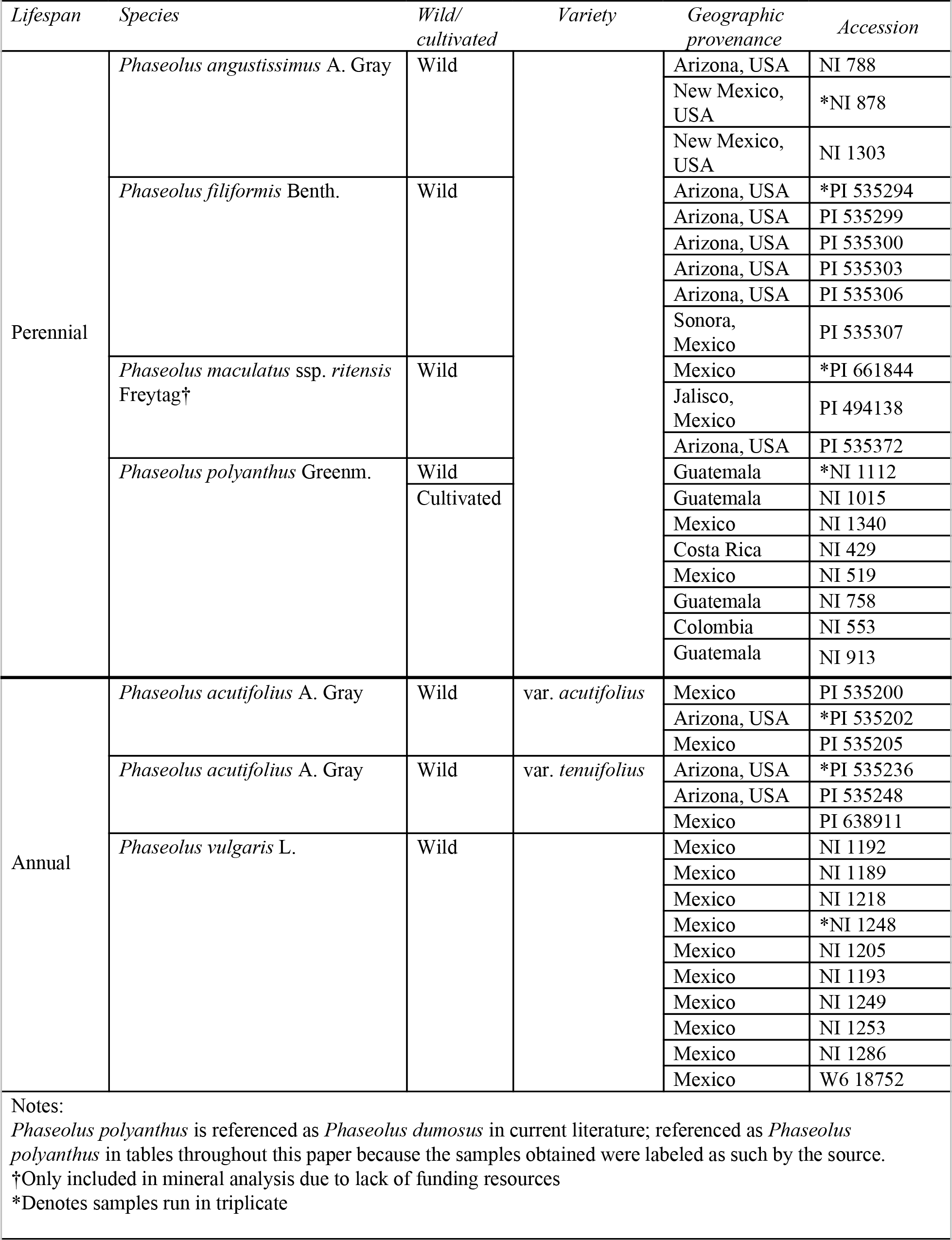
*Phaseolus* species analyzed with corresponding lifespan, cultivation status, geographic provenance, and accession number. Accession numbers that begin with ‘PI’ where obtained from the USDA and those that begin with ‘NI’ were obtained from Meise Botanic Garden of Belgium. *P. polyanthus* is referenced as *P. dumosus* in current literature; referenced as *P. polyanthus* in tables and throughout this paper because the samples obtained were labeled as such by the source.† denotes species only included in mineral analysis. * denotes samples run in triplicate.

Seeds were obtained from two sources: the United States Department of Agriculture (USDA) Western Regional PI Station (Pullman, WA) and the Meise Botanic Garden of Belgium and originated from Colombia, Mesoamerica (Costa Rica, Guatemala, and Mexico) and the southwest United States (Arizona and New Mexico) (Table 1). Seeds from the USDA were grown in a greenhouse for bulk seed production. Seeds from Belgium were obtained later than the USDA seeds, and consequently were unable to grow them in the greenhouse prior to processing; as a result, seeds from Belgium were processed directly. Differences in seed sources are accounted for in statistical analyses (see below). Seeds obtained from the USDA were germinated and moved to a hoop-house at the Missouri Botanical Garden (St. Louis, MO, USA) from May through August 2017. Plants were grown in 38-cell trays (cell diameter: 5.0 cm, cell depth: 12.5 cm) in Ball Professional Growing Mix. Fertilizer (150 ppm of 15-5-15 NPK) was applied five times in their first two months of growth. In August the plants were relocated to a greenhouse at Saint Louis University with supplemental indirect light and largely unregulated temperatures. They were watered twice per week. To minimize the effect of spatial heterogeneity, accession replicates were randomized among trays and trays were rotated several times throughout the duration of the bulking generation. Seeds were collected from three to ten accessions of each species from the greenhouse bulking generation.

Amino acid analyses were conducted on five of the six *Phaseolus* species (excluding *P. maculatus*) at the Proteomic and Mass Spectroscopy Facility at the Donald Danforth Plant Science Center (St. Louis, MO) using chemistry based on Waters’ AccQ•Tag derivatization method [59], [60]. The analysis quantified the concentration (pmoles/μl) of eight essential amino acids: histidine (His), isoleucine (Ile), leucine (Leu), lysine (Lys), methionine (Met), phenylalanine (Phe), threonine (Thr), valine (Val), and eight non-essential amino acids: alanine (Ala), arginine (Arg), aspartate (Asp), glutamine (Glu), glycine (Gly), proline (Pro), serine (Ser), and tyrosine (Tyr).

### Statistical analysis

For each ion, the mean and standard deviation was estimated. Data points with values greater than three standard deviations from the mean for each ion were removed. This resulted in the removal of 22 ion data points, including 17 from the same sample of a single accession of *P. polyanthus* (NI 553). For 20 of the 21 ions, one data point was removed, while two data points were removed for the remaining ion (Se). Preliminary analyses revealed that both ion and amino acid concentrations varied significantly by seed source (USDA *vs*. Meise Botanic Garden of Belgium). To control for this variation, we ran a linear model for each ion/amino acid that included only the fixed effect of seed source (SAS PROC GLM). From this model we extracted variation not explained by seed source (residuals) for use in subsequent models. To facilitate comparisons across ions and amino acids, residual data were standardized to a mean of zero and a standard deviation of one (SAS PROC STANDARD). For the seven accessions with biological replicates, the mean value was calculated and used for all downstream analyses.

Linear models were used to investigate fixed effects of lifespan and species nested within lifespan for each ion and amino acid (SAS PROC GLM). If a significant ‘species nested within lifespan’ term was present, we used contrast statements to test for pairwise differences between species within a lifespan to determine if annual species, perennial species, or both groups were variable, including a Bonferroni correction for multiple tests. To test for differences between wild and cultivated accessions and between varieties of the same species, separate fixed terms for these effects were also included in each model. Pearson correlation coefficients were estimated for all pairwise combinations of ions and pairwise combinations of amino acids (SAS PROC CORR). The correlation matrix was then used to cluster ions and amino acids into groups based on patterns of covariation using the rquery.cormat (graphType="heatmap") function in R [61]. To test for overall relationships between ion and amino acid concentrations, a principal components analysis (PCA; SAS PROC PRINCOMP) was first performed to reduce the dimensionality of the data. PCs that explained more than 5 percent of the variation were included in subsequent analyses, resulting in 6 axes of variation for ions (iPC1-6) and 2 axes of variation for amino acids (aPC1-2). Lastly, we estimated Pearson correlation coefficients between the ion and amino acid PCs (SAS PROC CORR). Data were analyzed in SAS and visualized using the ggplot2 R package [62], [63].

## RESULTS

Initial analyses determined that ion and amino acid concentrations differed based on seed source (USDA or Meise; Tables S1 and S2). Since the purpose of this study was to examine variation between lifespans and within and among species, we used statistics to account for variation due to seed source. As a result, in all subsequent analyses, we were able to compare *Phaseolus* nutritional content regardless of seed source.

### Differences among perennials and annuals in ion and amino acid concentrations

We assessed ion concentrations in two annual species (one of which included two varieties) and four perennial *Phaseolus* species and determined that nine of the 21 ions differed across species with different lifespans (Table 2). Specifically, calcium, copper, iron, magnesium, manganese, phosphorus, strontium, and sulfur were significantly lower in perennial species relative to annual species, while sodium was enriched for perennials (Figure 1 and Table 2). With respect to human nutrition, it is worth noting that the essential mineral nutrient, zinc, did not differ across annual and perennial species. Similarly, aluminium and arsenic, which in high concentrations pose a health risk, are equal across annuals and perennials.

**Table 2.**
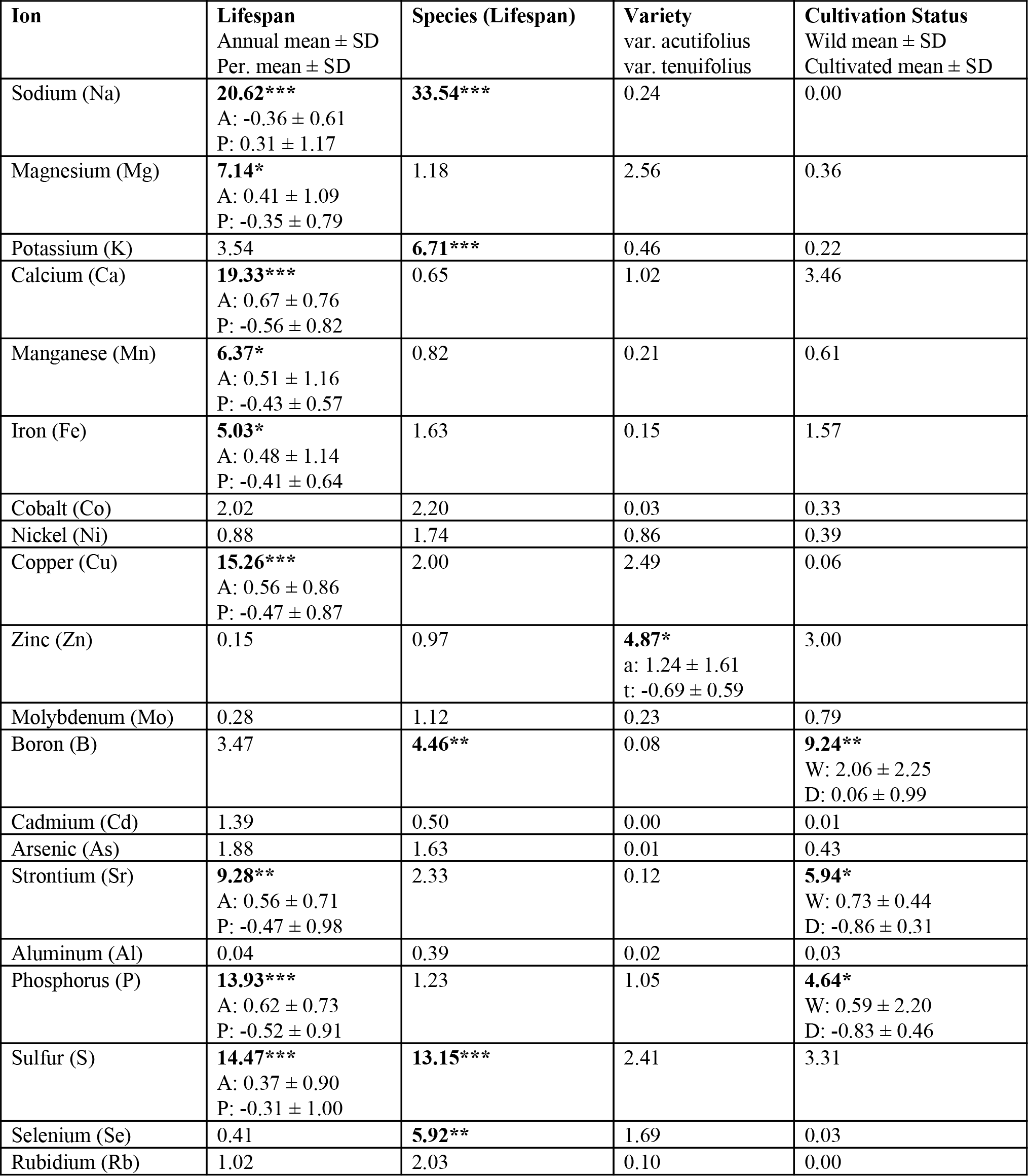
Results of linear models evaluating the effect of lifespan (annual *vs*. perennial), species nested within lifespan, variety and cultivation status on the concentration of 21 ions in whole seeds. All model terms were treated as fixed effects and *F*-values are reported. *P*-values: \**P* < 0.05, \*\**P* < 0.01, \*\*\**P* < 0.001; significant values are bolded. For significant lifespan, variety or cultivation status terms, means and standard deviations are shown.

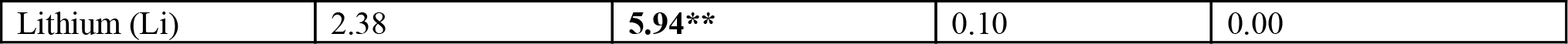

**Figure 1.**
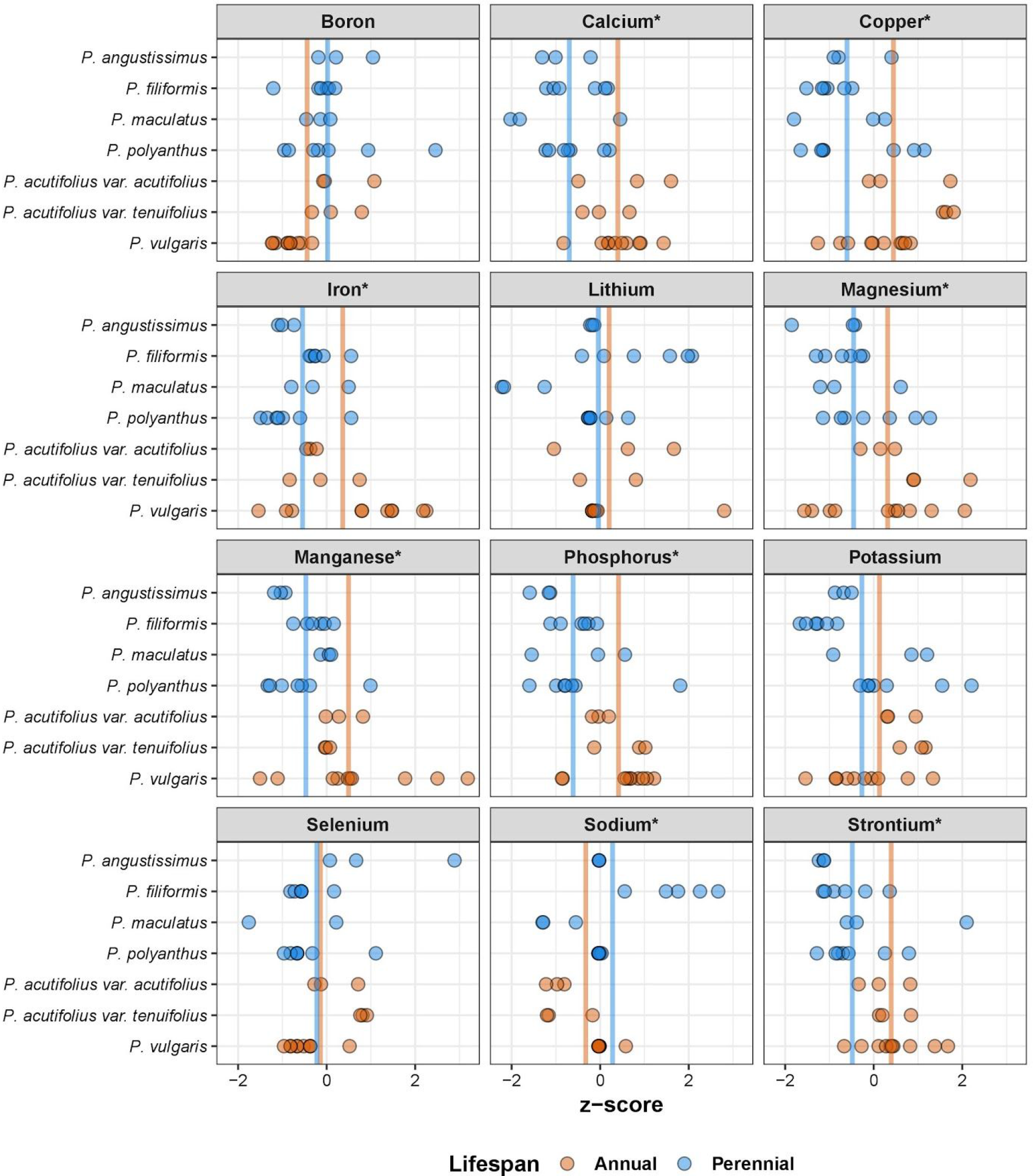
Concentrations (standardized z-scores) of 12 of the 21 ions quantified across perennial and annual *Phaseolus* species. Perennial species are shown in blue and annual species in orange. Each point represents a single accession. The intensity of the color increases with more overlapping data points. Vertical lines depict the mean of the perennial (blue) and annual (orange) species. * indicates ions that are significantly different (*P* < 0.05) between perennial and annual species. Ions included without an asterisk had significant variation among species *within* a lifespan; see Table 2 for a complete list of which ions had a significant lifespan and/or species effect.

In contrast to ions, amino acid concentrations generally did not vary between perennial and annual *Phaseolus* species. There were no significant differences based on lifespan for any eight of the essential amino acids, nor were there significant differences based on lifespan for six of the eight non-essential amino acids quantified in this study (Figures 2, S3 and Table 3). Two non-essential amino acids varied among lifespan: serine was significantly higher in perennial species compared to annuals, and tyrosine was significantly lower in perennials relative to annuals.

**Table 3.** Results of linear models evaluating the effect of lifespan (annual *vs*. perennial), species nested within lifespan, variety and cultivation status on the concentration of 16 amino acids in whole seeds. All model terms were treated as fixed effects and *F*-values are reported. *P*-values: \**P* < 0.05, \*\**P* < 0.01, \*\*\**P* < 0.001; significant values are bolded. For significant lifespan, variety or cultivation status terms, means and standard deviations are shown.

| Amino Acid | Lifespan<br>Annual mean ± SD<br>Per. mean ± SD | Species<br>(Lifespan) | Variety | Cultivation Status |
| --- | --- | --- | --- | --- |
| Histidine (His) | 0.06 | 1.60 | 1.91 | 0.45 |
| Serine (Ser) | **5.92\***<br>A: $-0.39 \pm 0.77$<br>P: $0.42 \pm 1.07$ | **3.97\*** | 0.76 | 2.03 |
| Arginine (Arg) | 0.01 | 0.32 | 1.83 | 0.03 |
| Glycine (Gly) | 0.06 | 0.37 | 1.98 | 0.36 |
| Aspartic acid (Asp) | 0.01 | 1.4 | 2.64 | 0.57 |
| Glutamic Acid (Glu) | 0.00 | 0.07 | 3.42 | 0.32 |
| Threonine (Thr) | 0.56 | **4.00\*** | 2.42 | 0.76 |
| Alanine (Ala) | 0.07 | 0.74 | 3.15 | 0.13 |
| Proline (Pro) | 0.75 | 0.19 | 2.38 | 0.60 |
| Lysine (Lys) | 3.73 | 1.56 | 2.58 | 0.28 |
| Tyrosine (Tyr) | **5.74\***<br>A: $0.44 \pm 0.87$<br>P: $-0.47 \pm 0.94$ | **3.11\*** | 0.92 | 1.66 |
| Methionine (Met) | 0.37 | 0.30 | 1.45 | 0.44 |
| Valine (Val) | 0.08 | 0.51 | 2.56 | 0.14 |
| Isoleucine (Ile) | 0.25 | 0.52 | 1.93 | 0.40 |
| Leucine (Leu) | 0.17 | 0.42 | 1.79 | 0.36 |
| Phenylalanine (Phe) | 0.75 | 0.82 | 1.86 | 0.41 |

**Figure 2.**
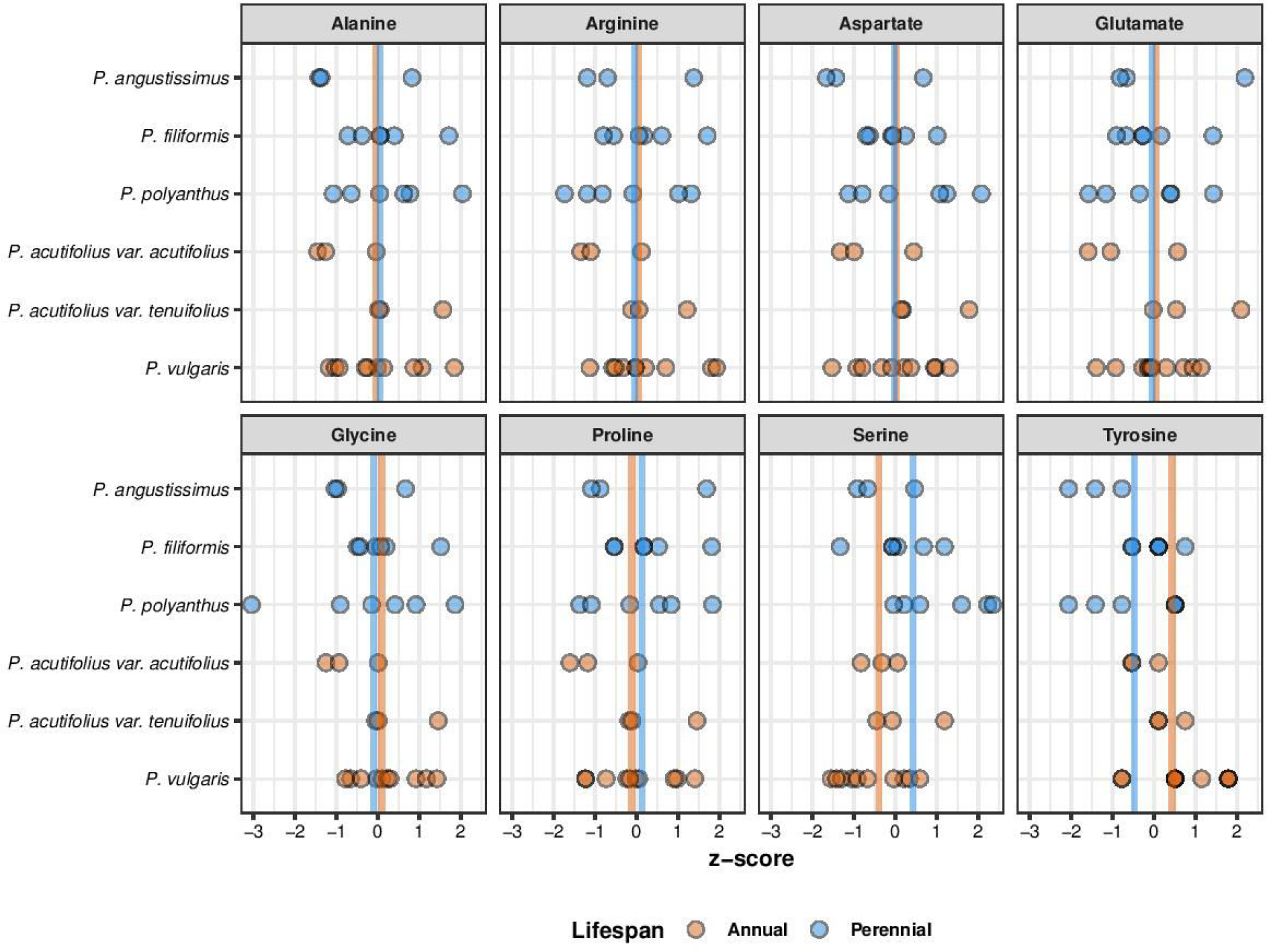
Concentrations (standardized z-scores) of eight essential amino acids quantified across perennial and annual *Phaseolus* species. Perennial species are shown in blue and annual species in orange. Each point represents a single accession. The intensity of the color increases with more overlapping data points. See Table 3 for a complete list of which amino acids had a significant lifespan and/or species effect.

### Variation within species in ion concentrations and amino acid concentrations

We observed high intraspecific variability of ion concentrations within and among accessions within a species (Figures 1, S1 and S2). For example, *P. angustissimus* accessions ranged from 56-69 ppm of iron and 42-90 ppm of zinc. *P. filiformis* accessions ranged from 59-91 ppm of iron and 35-97 ppm of zinc. Similarly, *P. maculatus* accessions contained 44-89 ppm of iron and 21-102 ppm of zinc. There was no statistically significant variation among species within lifespan category (annual or perennial) for the majority of ions (16 of 21). However, five ions (boron, potassium, selenium, sodium, and sulfur) differed among species within lifespan, for both annual species and perennial species (Figure 1 and Table 2). Case studies shed additional light on intraspecific variation in ion and amino acid concentrations. The first case study examined intraspecific variation in two varieties of *P. acutifolius*. Within *P. acutifolius*, ion concentrations were not variable across the two varieties, with the exception of zinc, which was higher in var. *acutifolius* relative to var. *tenuifolius* (Table 2). In the second case study we examined variation in ion concentrations and amino acids in wild vs. cultivated accessions of *P. polyanthus*. Three ions (boron, phosphorus and strontium) were significantly different between wild and cultivated accessions: concentrations of boron, phosphorus, and strontium were lower in cultivated accessions relative to wild accessions (Table 2).

For amino acids, we observed significant variation in one essential amino acid, threonine, and two non-essential amino acids, serine and tyrosine, within perennial species; however, none of the amino acids significantly differed between the two annual species (Table 3). For the case studies examining intraspecific variation and variation in cultivated vs. wild accessions, no differences in amino acids were detected (Table 3).

### Correlations among ions and amino acids

We evaluated the relationships among nutrients by performing pairwise correlations within the 21 ions and within the 16 amino acids (Figures 3 and 4). Of the 210 possible pairwise ion correlation combinations, 70% were non-significant (147/210). We detected 63 significant correlations among ion concentrations (30%), and of those, there were more positive than negative associations (49 vs. 14), with correlation coefficients (r) ranging from −0.57 to 0.77. In contrast, of the 120 possible pairwise amino acid correlation combinations, 94% (113/120) were significantly correlated. The majority of the among amino acid correlations were positive (85%, 102/120; Figure 4), while the remaining 11 were negative. All of the negative correlations were between methionine and other amino acids.

**Figure 3.**
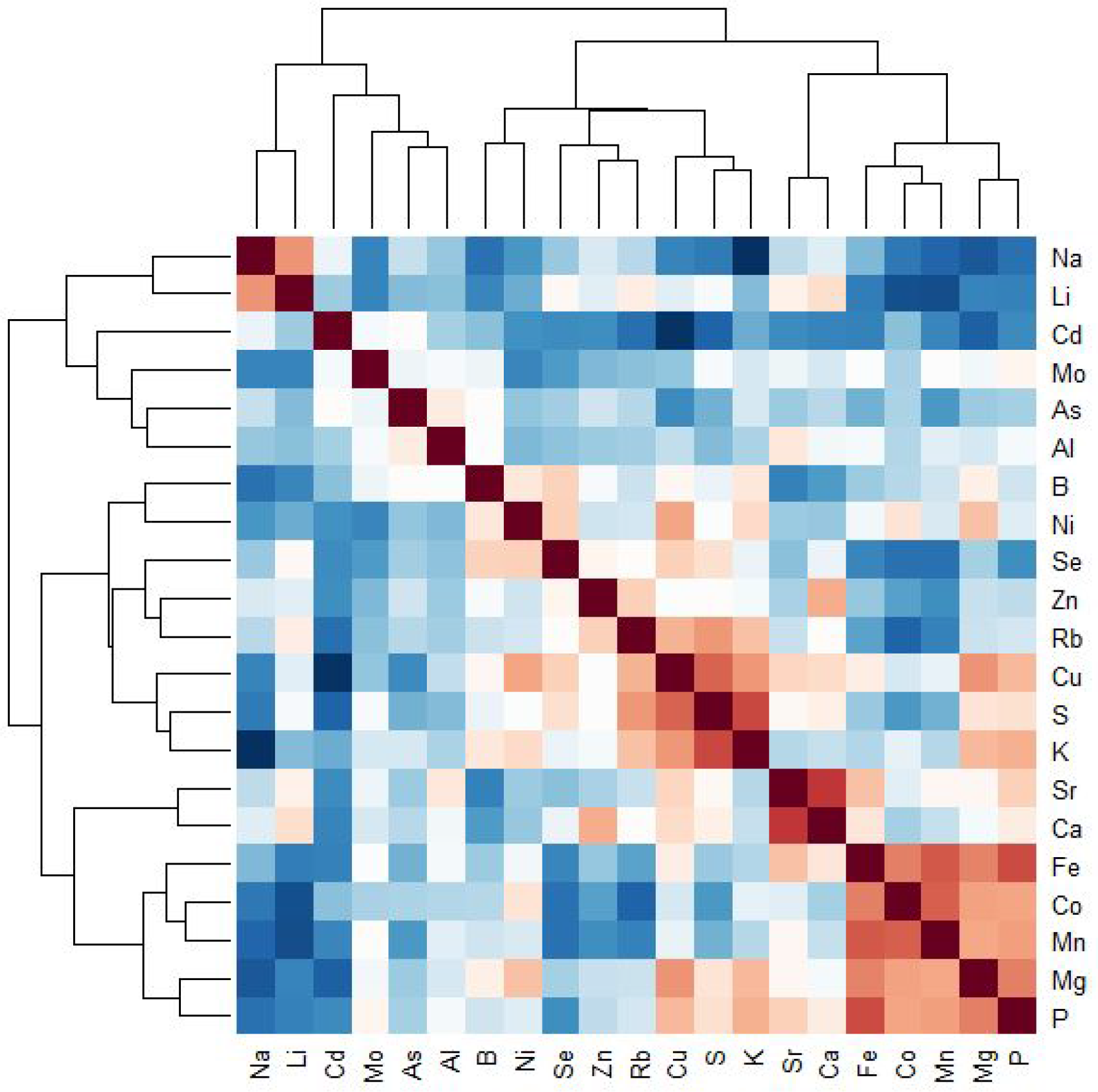
Cluster analysis, based on patterns of covariation, for the 21 ions quantified across perennial and annual *Phaseolus* species. Correlation coefficients range from −1.0 (dark blue) to 1.0 (dark red).

**Figure 4.**
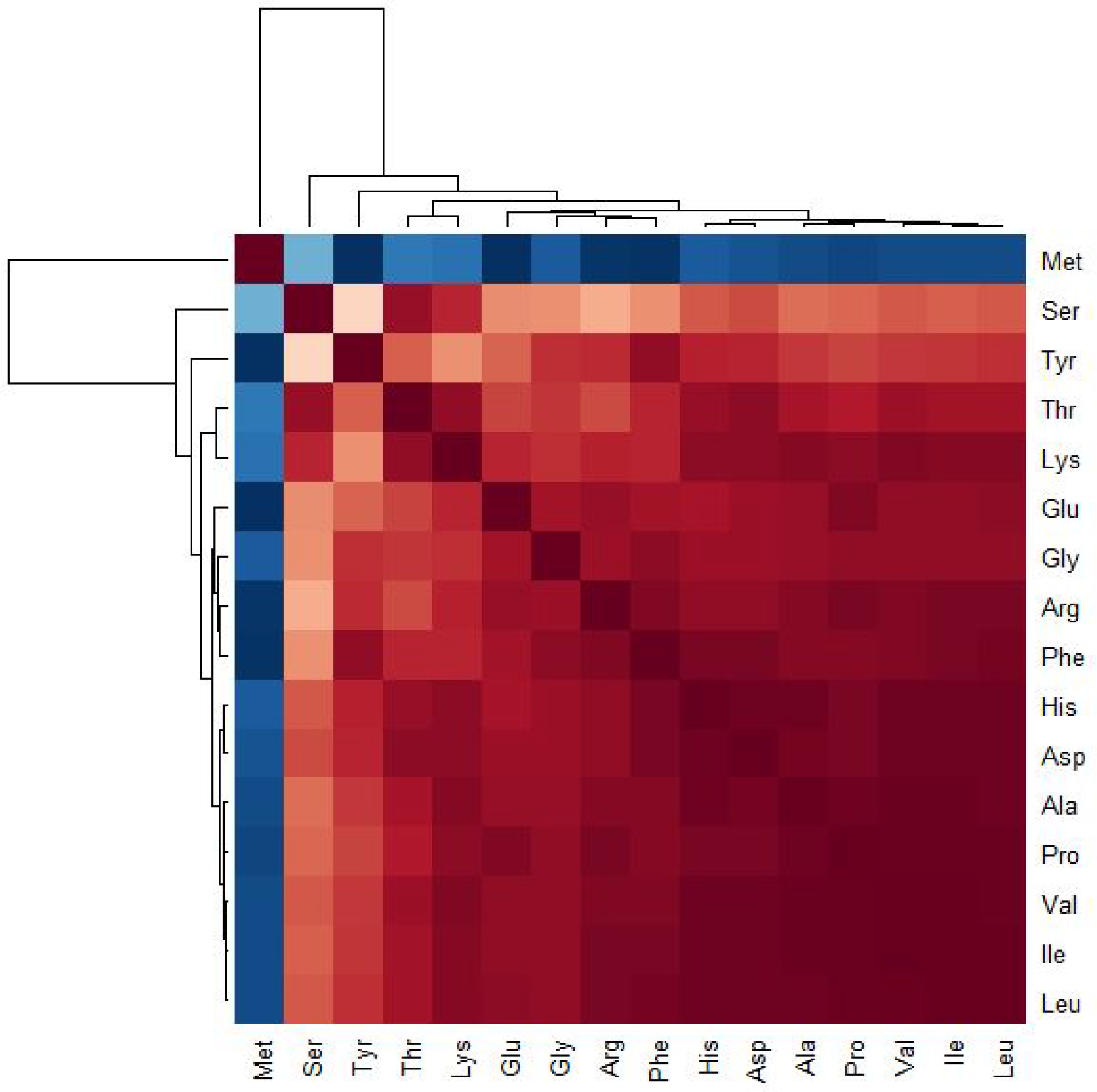
Cluster analysis, based on patterns of covariation, for the 16 amino acids quantified across perennial and annual *Phaseolus* species. Correlation coefficients range from −1.0 (dark blue) to 1.0 (dark red).

To further explore relationships among ions and among amino acids beyond single bivariate correlations, we used a clustering approach to determine groupings based on overall correlation structure (i.e., are there groups of ions that have similar relationships with all other ions). Clustering algorithms applied to ion concentrations identified two large groups of ions based on their covariation patterns, one with six ions (Na, Li, Cd, Mo, As, and Al) and a second with the remaining 15 ions (Figure 3). Within the second group of 15 ions, some sub-clustering was apparent. For example, one subcluster consisted of Fe, Co, Mn, Mg, and P; a second included Sr and Ca; a third subcluster included S and K (Figure 3). The amino acids formed two clusters, the first consisting of only methionine and the second consisting of the remaining 15 amino acids (Figure 4). To test for an effect of lifespan and species on multi-element ion and amino acid clusters, we generated six principal components for ions (iPC) and two amino acid principal components (aPCs; Table S3 and S4). Principal component analysis on the ions and amino acids yielded similar results to the individual ion and amino acid analyses; both lifespan and species were variable for some PCs (Table S5). To test for broad relationships between ion and amino acid concentrations, we tested for associations between PCs rather than individual elements. Of the 12 potential bivariate correlations among the ion and amino acid PCs, only iPC2 and aPC2 were significantly positively correlated.

## DISCUSSION

Increasing pressure on agriculture has focused attention on improving crop yield and nutrition. In addition, environmental considerations are driving development of productive systems that also reduce negative impacts on soil, water, and biodiversity resources. Agroforestry, which combines agriculture and forestry, offers one option for how agricultural systems might meet nutritional needs while also improving ecological sustainability. In recent years attention has focused in part on the possible development of perennial, herbaceous crops [11], [13], [14], [64]. Expanding development of species such as wild, perennial legumes, into an agroforestry system offers opportunities to enhance food production and possibly reduce the risk of nutrient deficiencies [6], [7], [65], [66]. In this study, we explored differences in nutrient content among perennial and annual wild species of the common bean genus *Phaseolus*, an important first step towards identifying *Phaseolus* species that might be good candidates for pre-breeding and/or inclusion in agroforestry and other agricultural systems.

### Comparisons of ion and amino acid concentrations in annual and perennial Phaseolus

As ecological intensification of agriculture prioritizes exploration of perennial crops, one goal is to equal or enhance nutritional value relative to existing systems. Our results suggest that relative to annual *Phaseolus* species, perennials provide comparable protein quality, zinc levels, and are equally limited in some toxic compounds (e.g. arsenic). Further, amino acid concentrations between perennial and annual *Phaseolus* species were comparable, with the exception of two non-essential amino acids, serine and tyrosine. These data, among other studies [19], [34], [48], [52], [67], [68] suggest that, from a nutritional perspective, perennial relatives of traditional annual crops like *Phaseolus* and others warrant additional evaluation. Our work expands what modest nutrient information was known for perennial *Phaseolus* species and statistically compares to closely related annual species.

While perennial and annual *Phaseolus* species share essential nutritional characteristics, there are some differences among accessions across lifespans. Relative to annuals, perennial *Phaseolus* have lower levels of some essential mineral nutrients including calcium (41%), iron (22%), and magnesium (13%). In contrast, a previous study [34] found that perennial wheat species exceeded annual species in some essential mineral nutrients. In a perennial by annual hybrid (*Thinopyrum elongatum* x *Triticum aestivum)* grain were 24, 32 and 33% higher in iron, magnesium and zinc, respectively, compared to annual parent species, *Triticum aestivum* L. Increased ion absorption in the progeny was attributed to deeper root systems [34]. Thus, we would expect improved ion scavenging by the perennial species in comparison to annuals. However, many of our perennial seed samples were harvested from relatively young plants grown in small pots, restricting the root systems and resulting in root structures comparable to annual species. Additional comparative analyses of closely related perennial and annual species, either in large pots or ideally in the field, are needed to infer general patterns with regards to nutritional composition and lifespan.

The application of high throughput ionomics and proteomics pipelines represents an important opportunity to explore natural variation in plant species for agricultural improvement, but also to ask fundamental questions in plant biology. Work presented here identified differences in ion concentrations among wild perennial and annual *Phaseolus* species, focusing attention on the broader question of how perennial and annual species differ from one another. One reason why perennial and annual species might differ in their nutritional compositions is that perennial and annual species allocate resources differently due to differences in their lifecycle demands [14], [69], [70]. For example, due to their dependence on sexual reproduction for survival, annual species tend to allocate a greater proportion of their biomass to seeds than herbaceous perennials [71], [72]. The result of this differential allocation is that, in any given year, perennial species tend to produce fewer seeds relative to annual species [72]. One potential effect of seed allocation differences between perennial and annual plants is the dilution effect, which results in a reduction in nutrient (ion and protein) concentration as grain size increases. However, it is unclear if there is a consistent difference in seed size among perennial and annual congeners [72]; Herron unpublished data). Annual root systems are also characterized by high nitrogen concentration and root specific length, which implies higher resource acquisition from the soil, whereas perennials maintain denser, more durable roots which may grow to greater depths and new nutrient pools [73]. Another potential factor shaping variation in ion concentrations in seeds is the location of the seed and seed pod on the plant. Plant parts in proximity to seeds (e.g., leaves, pod walls, etc.) may preferentially partition ion contents to nearby seeds [74]. Position of inflorescences along a plant’s stem has been shown to impact nutrient allocation in soybean seeds [30] and grapevine leaves [28]. Future work is needed to comprehensively sample *Phaseolus* seeds from throughout the plant in order to determine the effect of seed position on nutrient concentrations in this genus.

### Natural variation in mineral and amino acid concentrations within six wild Phaseolus species

Ion concentrations and amino acids vary across species examined, indicating that natural variation exists in nutritional components of plants, and that this variation may be important for future breeding efforts. In addition to detecting variation among lifespan (annual and perennial species), we observed variation in ion levels within species. Differences in ion concentrations tended to be ion-specific, with some ions exhibiting more extensive variation than others (e.g., iron levels varied significantly, whereas cadmium did not). Some ions appeared to vary independently of one another (see discussion below). In addition, amino acid concentrations were variable among *Phaseolus* species as well; however, accessions with high or low concentrations of one amino acid tended to have high or low levels across all amino acids. Thus, accessions with high amino acid concentrations may warrant further attention when considering artificial selection for nutrient quality [75]. Our results are supported by previous work identifying variability in nutrient values between accessions within a species [18], [20], [76]–[79]. Intraspecific variability in wild *Phaseolus* species is likely due to a combination of genetic and environmental factors [18], [20], [77], [78]; but regardless represents a rich source of variation that warrants exploration and possibly pre-breeding.

Two cases studies shed additional light on variation in ion and amino acid concentrations. First, we compared two varieties of annual *P. acutifolius,* var. *acutifolius* and var. *tenuifolius*. These two varieties are differentiated by leaf morphology and occupy similar geographic areas [80]. Recent studies show that these varieties confer drought and frost tolerance when crossed with cultivated *P. vulgaris* [81]. We observed significant differences in zinc levels between the two varieties of *P. acutifolius*, however, all other essential nutrients and most ions were comparable across varieties. Our results are consistent with previous work which identified similar ion and amino acid concentrations across varieties adapted to similar environments [19], [82], [83]. A second case study investigated differences between wild and cultivated species of *P. polyanthus.* Similar to naturally evolved varieties of *P. acutifolius*, we found little variation between wild and artificially selected (cultivated) accessions of *P. polyanthus*. The only significantly different essential ion between wild and cultivated *P. polyanthus* was phosphorus, which was higher in the wild species. They also deviated in two non-essential ions, boron and strontium (Table 2). Likewise, other studies of *Phaseolus* have found wild accessions of *P. vulgaris* to have higher protein and elevated levels of some mineral nutrients compared to cultivated accessions [84], [85]. These data further underscore the existence and importance of natural variation in ion and amino acid concentration as a resource to be conserved and integrated into breeding efforts.

### Correlations in mineral and amino acid concentrations across Phaseolus species

Correlations among traits can facilitate or impede artificial selection depending on the direction and strength of the correlation. With respect to breeding this can be problematic when selection on increased concentration of a nutratively beneficial ion also increases the concentration of a potentially harmful ion. For ion concentrations, we found that 70% of bivariate trait correlations were non-significant suggesting that breeders may be able to optimize the concentrations of ions fairly independently and thus generate a final product rich in beneficial ions (e.g., iron and zinc) but low in harmful ions (e.g. arsenic).

The remaining ion concentration bivariate trait correlations (30%), as well as the amino acid concentrations, co-varied with one another. For example, we identified clusters or groups of ions that covaried with other ions, a pattern that has been detected in a wide-range of plant taxa, including *Arabidopsis thaliana*, *Brassica napus*, *Glycine max*, and *Phaseolus vulgaris* across multiple tissues [20], [25], [86]–[88]. In one study in the model plant, *A. thaliana*, genetic studies demonstrated that the concentrations of multiple ions are controlled by the same genomic region [25], [89]. These genomic regions often contain genes involved in ion transport suggesting that multiple ions may be absorbed from the soil and/or moved through the plant by the same mechanism [86], [90]. In our study, we find correlations that are both similar (e.g., calcium & strontium; [91], [92] and different (e.g., copper, zinc, and nickel; [93] than previous studies in leaves [94]. Collectively, the growing body of literature on ion covariation suggests that ion concentrations are highly dynamic and therefore it will be important for breeders to determine the ion variation and covariation for each species (and possibly across multiple geographic locations). However, the promise of agricultural crops with optimization of multiple ions is obtainable; recently, iron and zinc have been successfully co-selected for increased concentrations in wheat species [66].

Patterns of covariation were far less variable for the amino acids quantified. Most (15/16) amino acids clustered into one large group, and within group bivariate correlations were always positive, generally significant, and quite strong. Strong correlations among amino acids are less of a concern to breeders because, unlike ions, none of the amino acids measured are potentially harmful in normal quantities. Therefore, artificial selection for increased concentrations of any amino acids will likely result in increased concentrations of all other amino acids, with the exception of the essential amino acid methionine. Methionine was significantly negatively correlated with most other amino acids. Therefore, it is possible that breeding for high levels of 15 amino acids in *Phaseolus* species may ultimately result in a reduction of methionine. A previous study on correlations among amino acids within proteins indeed found a negative correlation between methionine and eight of the 15 other amino acids included here [95]. However, *Phaseolus* seeds tend to be a poor source of sulfur-containing amino acids (e.g. methionine and cysteine), therefore a healthy diet including other complimentary plant sources (e.g. oats, soybean) or animal products will supply sufficient methionine [77], [96], [97].

### Future directions

Developing crops to meet nutritional needs of a growing population and at the same time enhancing ecological sustainability of agricultural systems is a core goal of current breeding programs. As new candidates are considered for agroforestry and other forms of agriculture, nutritional quality of candidate species should be assessed in order to maximize nutrient output of sustainable food crops. Our work as well as other recent studies indicates that there are similarities among perennial and annual *Phaseolus* species in nutrition components, and that there is variability among accessions within a species, providing the opportunity to select for essential nutrients by plant breeders [75], [76], [98]. Future expansion could include surveying a wider range of both wild species and accessions to understand nutrient uptake dynamics in a wider phylogenetic and geographic context. Moreover, within-plant variation is important from an evolutionary and nutritional standpoint. To better understand these processes, it is important to characterize the pool of ions/amino acids available to the maternal plant from the soil, the overall levels in the maternal plant, and the patterns of allocation to developing beans over space and time. Among perennial species, root system characteristics (e.g., depth, fibrous vs. taproot) may be compared to establish ion uptake mechanisms. In addition, anti-nutrient analysis, organoleptic and toxicology studies would contribute to a more comprehensive understanding of the human nutrition applications of wild, perennial beans.

Future studies should consider other perennial species in general, and within *Phaseolus* future next steps might include assessment of variation within and across clades examined here, such as *P. coccineus* and *P. polystachios,* respectively. Proteomic and ionomic analyses should also be replicated for all accessions in this study to test for repeatability of concentration levels. It is only by capturing a broad phylogenetic range and ensuring that accessions have high concentrations in field conditions that we will be able to identify accessions with optimal nutrient quality. Ultimately, this work can lead to the selection of accessions for use in pre-breeding programs where nutrient quality should be considered in conjunction with other agriculturally important factors. These nutritionally rich, perennial herbaceous legume species could complement existing agriculture and agroforestry systems, diversifying the ecosystem and elevating nutrient output.

## CONCLUSION

Comparing annuals and perennials helps develop a better understanding of how the integration of perennial crops into agriculture may impact the nutrient pool of our food system. Results from this study indicate there are nutritional differences at the lifespan and species level. Ionomics data revealed annual species exceed perennials in calcium, iron, magnesium, manganese, phosphorus and selenium, while zinc levels were comparable. However, virtually no differences were observed among amino acid profiles, suggesting protein quality of perennial species are comparable to annual species. The range in intraspecific variation among individual accessions of the same species further demonstrates the complexity of nutrient allocation and the need for more research assessing the variation in nutrient content in underutilized, perennial crops. This variation speaks to the opportunity to artificially select for nutritionally important compounds within ecologically important species. This study provides a snapshot of what perennial crops may potentiate, as there are thousands of perennial legume species with very little data regarding nutritive compounds. Further studies are needed to replicate analyses of the species within this study and to explore the thousands of underutilized herbaceous, perennial legume species.

## AUTHOR CONTRIBUTIONS

conceptualization, H.S., K.E., L.L and A.M.; methodology, H.S., K.E., S.H., L.L and A.M.; statistical analysis, M.R.; plant growth and sample preparation, H.S., S.H.; data curation, H.S. and M.R.; writing—original draft preparation, H.S., M.R., and A.M; writing—review and editing, H.S., K.E., S.H., L.L, Z.M., M.R. and A.M.; data visualization, M.R. and Z.M.; supervision, K.E., L.L. and A.M..; project administration, H.S. and S.H.; funding acquisition, H.S., K.E., L.L and A.M.

## ACKNOWLEDGEMENTS

Special thanks to members of the Miller Lab for their critical review of the manuscript. To the following from the Missouri Botanical Garden: Derek Lyle (for providing the space), and Justin Lee, Joshua Higgins, Claudia Ciotir, and Summer Sherrod for their assistance in plant care. To the following from Saint Louis University: Emma Frawley and William Shoenberger for their assistance in plant care. To the following from Baldwin Wallace University: Dr. Raymond Shively and Dr. Diana Barko for their contributions with chemical equations. In addition, the Meise Botanic Garden and United States Department of Agriculture for providing seed material and the Proteomics & Mass Spectrometry Facility and Baxter Ionomics Facility at the Donald Danforth Plant Science Center for conducting the data generation.

## SUPPLEMENTARY MATERIALS

**Figure S1**. Concentrations (standardized z-scores) for the seven accessions with replicate data points all ions measured for one accession of each *Phaseolus* species (and variety). Perennial species are shown in blue and annual species in orange.

**Figure S2**. Concentrations (standardized z-scores) of nine ions that were not variable within or across perennial and annual *Phaseolus* species. Perennial species are shown in blue and annual species in orange. Each point represents a single accession. The intensity of the color increases with more overlapping data points. Vertical lines depict the mean of the perennial (blue) and annual (orange) species.

**Figure S3**. Concentrations of 8 non-essential amino acids across perennial and *Phaseolus* species. Perennial species are shown in blue and annual species in orange. The intensity of the color increases with more overlapping data points. Vertical lines depict the mean of the perennial (blue) and annual (orange) species.

**Figure S4**. Clustering analysis, based on patterns of covariation, for the 6 ion (iPC1-6) and 2 amino acid (aPC1-2) PCs generated. Correlation coefficients range from −1.0 (dark blue) to 1.0 (dark red).

**Table S1**. Preliminary linear models testing for an effect of seed origin (USDA vs. Meise Botanic Garden of Belgium) on the 21 ions measures across perennial and annual *Phaseolus* species. All model terms were treated as fixed effects and *F*-values are reported. *P*-values: \**P* < 0.05, \*\**P* < 0.01, \*\*\**P* < 0.001; significant values are bolded.

**Table S2**. Preliminary linear models testing for an effect of seed origin (USDA vs. Meise Botanic Garden of Belgium) on the 16 amino acids measures across perennial and annual *Phaseolus* species. All model terms were treated as fixed effects and *F*-values are reported. *P*-values: \**P* < 0.05, \*\**P* < 0.01, \*\*\**P* < 0.001; significant values are bolded.

**Table S3**. Loadings for the 21 ions measured onto the first 6 PC axes (iPC1-6).

**Table S4**. Loadings for the 16 amino acids measured onto the two 6 PC axes (aPC1-2).

**Table S5.** Results of general linear models evaluating the effect of lifespan (annual vs. perennial), species nested within lifespan, variety and cultivation status on the top 6 ion PCs and top 2 Amino Acid PCs. Proportion variance explained (PVE) is reported for each PC; only PCs that explained greater than 5 percent are shown. All model terms were treated as fixed effects and *F*-values are reported. *P*-values: \**P* < 0.05, \*\**P* < 0.01, \*\*\**P* < 0.001; significant values are bolded.

